# Integrative genomic approaches to study the barley-*Pyrenophora* teres interaction

**DOI:** 10.64898/2026.02.04.703911

**Authors:** Buddhika A. Dahanayaka, Lisle Snyman, Pavansai Bathini, Manaswini Sandiri, Sadegh Balotf, Anke Martin

## Abstract

*Pyrenophora teres* f. *teres* (Ptt), the causal agent of net-form net blotch in barley, was studied using a bi-parental mapping population (Pop1) of 305 isolates derived from a cross between two isolates with contrasting virulence on barley cultivars Skiff and Prior. QTL analysis identified virulence loci on chromosomes (Chr) 3 and 10 for Skiff, and on Chr 1, 4, and 5 for Prior. Major QTL on Chr 3 and 5 explained 24% and 40% of phenotypic variation, respectively. A second population (Pop2) was developed by crossing two Pop1 isolates, one carrying major QTL on Chr 3 and 5 and one avirulent. Isolates from Pop2 with single QTL were phenotyped across a Prior/Skiff recombinant inbred line population to identify corresponding host susceptibility/resistance loci. Skiff virulence QTL on Chr 3 corresponded to barley Chr 3H and 6H, while Prior virulence QTL on Chr 5 mapped to Chr 6H. RNA expression analysis of virulent and avirulent Pop2 isolates identified five candidate genes linked to the Chr 5 QTL, including two predicted effectors. These findings suggest both gene-for-gene and inverse gene-for-gene interactions in the Ptt–barley pathosystem and advance the understanding of molecular mechanisms underlying host-pathogen specificity.

## Introduction

Successful infection of a host plant by a pathogen is the result of a series of molecular events occurring during the invasion and colonization of the host plant. The ability of a host plant to fend off pathogenic attacks depends on its innate and adaptive immunity, and how well these immune systems can overcome the pathogen’s effector molecules, proteins which pathogens use to infect the host. The initial innate immune response occurs through host plant detection of pathogen-associated molecular patterns (PAMPs) by pattern recognition receptors (PRRs), whereby general immunity is induced (De Wit et al. 1997; Jones & Dangl 2006; De Wit et al. 2009). This process is known as PAMP-triggered immunity (PTI).

A number of genomic regions in the pathogen and in the host encode the components of the host immune system and the pathogen effector molecules, respectively. The expression of these genetic regions in host-pathogen interactions can result in qualitative or quantitative disease responses. Initially, the host-pathogen interaction was suggested to be qualitatively inherited or following a gene-for-gene interaction (Flor 1956). The gene-for-gene interaction describes instances where host plants possess resistance genes specific to a corresponding avirulence (*Avr*) gene in the pathogen, which triggers an immune response to prevent infection. These R genes encode proteins that can recognize the effector molecules produced by the *Avr* genes of the pathogen, typically leading to the activation of plant defence responses. The gene-for-gene interaction describes the next layer of plant-pathogen interactions, whereby pathogens evolve *Avr* genes encoding effectors that can suppress PTI. These effectors enable the pathogen to invade the host plant and, when the host plant possesses corresponding R genes, activate effector-triggered immunity (ETI) and induce stronger, hypersensitive immune responses to the pathogen (De Wit et al. 2009). This leads to a co-evolutionary arms race between the pathogen and the host plants, where the pathogen adapts to the R gene by either losing the *Avr* gene, mutating its effectors, or gaining novel effectors to suppress ETI by different means (De Wit et al. 1997; Jones & Dangl 2006; De Wit et al. 2009). With the identification of such effectors, it was proposed that the host-pathogen interaction of necrotrophic fungi does not necessarily follow the gene-for-gene model, or qualitative inheritance, but instead conforms to a process of both quantitative and qualitative inheritance (Friesen et al. 2008).

Net blotch is an economically important foliar disease of barley world-wide, caused by the necrotrophic ascomycetous fungus *Pyrenophora teres*. The pathogen exists as two forms, *P. teres f. teres* (*Ptt*) and *P. teres f. maculata* (*Ptm*), which cause net-form net blotch (NFNB) and spot-form net blotch (SFNB) symptoms in barley (*Hordeum vulgare* L.), respectively (Smedegård-Petersen 1971). The sexual reproduction of *P. teres* is regulated by a single mating type locus (*MAT1*). The mating type locus of *P. teres* exists as two alternative forms or idiomorphs, i.e. *MAT1-1* and *MAT1-2*. *P. teres* is a heterothallic fungus and for successful sexual reproduction, two compatible individuals of opposite mating types are required (McDonald 1963). The heterothallism of *P. teres* allows us to examine the sexual recombination occurring in the pathogen and to understand the barley-*P. teres* pathosystem.

To date, six bi-parental mapping studies have been conducted to discover genomic regions accounting for avirulence/virulence of *Ptt/Ptt* populations (Weiland et al. 1999; Beattie et al. 2007; Lai et al. 2007; Shjerve et al. 2014; Koladia et al. 2017; Martin et al. 2020b) and one *Ptt/Ptm* hybrid mapping population (Dahanayaka et al. 2022). A genome-wide association study (GWAS) (Martin et al. 2020b) and a multi-parental nested associated mapping (Dahanayaka & Martin 2023) study have also been conducted to identify genomic regions responsible for the virulence in *Ptt*. These mapping studies have identified QTL across all 12 chromosomes of the *P. teres* genome that contribute to virulence against a range of globally grown barley cultivars such as Beecher, Harbin, Kombar, Rika and Skiff, demonstrating the genetic complexity of the *P. teres*-barley pathosystem.

The genetic basis of barley resistance to *P. teres f. teres* has been studied for nearly a century, with quantitative inheritance first demonstrated by Geschele in 1928 (Geschele 1928). Several major loci have since been identified which include *Rpt1*, *Rpt2*, *Rpt3*, *Rpt5/Spt1*, and *Rpt7*, found on chromosome (Chr) 3H, 1H, 2H, 6H and 4H, respectively. Typically, for each of these loci, several QTL have been identified over multiple studies that have since been condensed into a single locus. For example, the first resistance locus *Pt1* was discovered in the Tifang × Atlas cross and was later accompanied by *Pt2* and *Pt3* (Mode and Schaller 1958). A separate locus, *Pt,,a*, was identified for resistance to an Australian isolate (Graner et al. 1996). Over time, due to overlapping resistance traits and mapping studies, these loci were consolidated into a single locus, *Rpt1*, on Chr 3H (Bockelman et al. 1977) along with subsequently identified loci *QRpts3L* and *QNFNBSLR.Al/S-3H* (Raman et al., 2003 and Lehmensiek et al., 2007). Within this locus, there have been two alleles identified, *Rpt1.a* and *Rpt1.b* (Bockelman et al. 1977).

Single QTL regions identified for *P. teres* interaction can be associated with both resistance and susceptible interactions. Well-known *Rpt5/Spt1* locus on Chr 6H, which was previously classified as *Pt_a_*, is a complex and highly important locus in the *P. teres* f. *teres*-barley interaction (Clare et al. 2020). This complexity stems from the existence of both dominant resistance and susceptibility alleles across different barley lines. Currently three alleles are described within this locus. Allele *Rpt5.f*, identified from CI5791/CI9819 (Manninen et al. 2006), is written with a capital “R” indicating dominant resistance as opposed to alleles *rpt5.k* and *rpt5.r*, identified from Kombar and Rika (Abu Qamar et al. 2008), which are lower case “r” to denote recessive resistance (Franckowiak and Platz, 2013). However, as noted above, the *Rpt5* locus is also classified as *Spt1*, due to the designation of this locus as susceptibility to *P. teres* (Richards et al., 2016). Since this designation, the two recessive resistant genes have also been reclassified as *Spt1.r* (previously *rpt1.k*) and *Spt1.k* (previously *rpt1.r*) (Clare et al. 2020). It is currently unclear whether *Spt1.r* and *Spt1.k* are two closely linked genes or a single gene with divergent allelic series where the interaction with *P. teres* is dependent on different effectors (Richards et al., 2016).

Above mentioned studies explain the complexity of the barley-*P. teres* pathosystem. Investigating both the QTL associated with the pathogen and the corresponding host responses will facilitate the identification of key genes and genomic regions involved in this interaction. To elucidate the genetic basis of plant-pathogen interaction, the current study aimed to 1. identify virulent/avirulent QTL using a bi-parental mapping population developed by crossing *Ptt* isolates that are virulent and avirulent on barley cultivars Skiff and Prior; 2. identify the corresponding resistance/susceptibility QTL in barley cultivars Skiff and Prior 3. recognise the putative genes associated with barley-*Ptt* pathosystem via integrated genomic approaches.

## Materials and Methods

### Pathogen isolates and populations

A bi-parental mapping population of 325 isolates was developed by crossing naturally occurring *Ptt* isolates NB81 (*Ptt: MAT1-1*) and HRS09127 (*Ptt: MAT1-2*) (Pop1) (Dahanayaka&Martin 2023). The isolate NB81 has been reported to be virulent on barley cultivar Prior and avirulent on Skiff, while HRS09127 is avirulent on Prior and virulent on Skiff, respectively (Fowler 2018). After QTL analysis of Pop1, individual Pop1_451 possessing the QTL for Prior virulence, based on its genotype and phenotype, was crossed with an avirulent individual (Pop1_155) from the same population to create the bi-parental mapping population Pop2 consisting of 88 *Ptt* isolates (Figure. 1, Table 1). Pop2 was created to enable the selection of progeny which only have one QTL. Bi-parental *Ptt* population Pop3 (*n* = 89) was developed by crossing a *Ptt* individual of the NB29/HRS09122 population (Martin et al. 2020), associated with virulence on barley cultivar Skiff, with isolate HRS14051 avirulent on Skiff. All the fungal crosses were developed as described in Dahanayaka et al. (2022).

**Figure 1.**
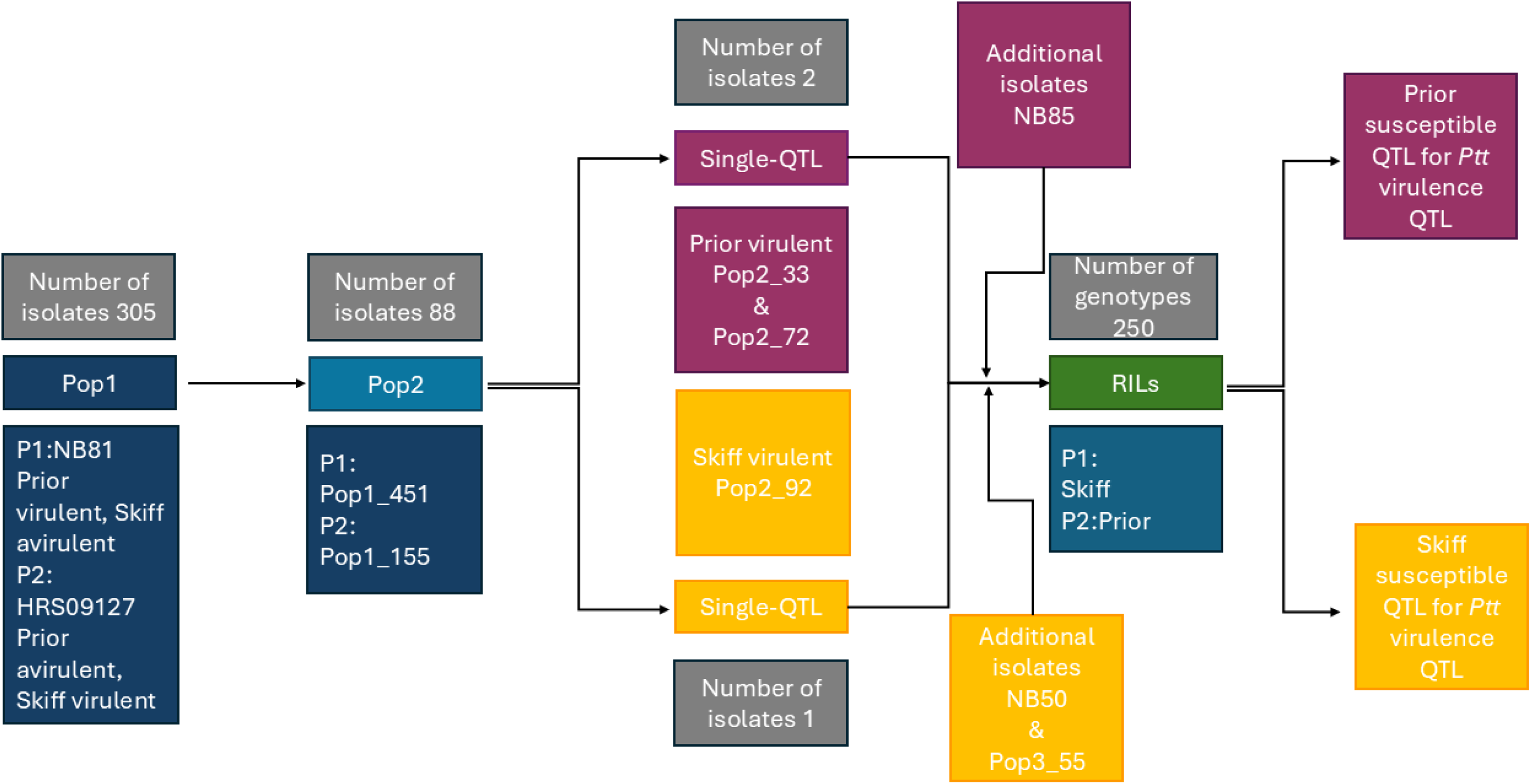
Flow-chart of *Pyrenophora teres* f. *teres* mapping populations used in this study for the identification of virulence genes associated with the Skiff and Prior barley varieties, and barley Skiff/Prior recombinant inbred line population used in the quantitative trait loci mapping of *Ptt* virulences.

**Table 1.** Information of fungal bi-parental mapping populations

| Cross ID | Parent 1 | Mating type | Skiff | Prior | Parent 2 | Mating type | Skiff | Prior | Total number of isolates |
| --- | --- | --- | --- | --- | --- | --- | --- | --- | --- |
| Pop1 | NB81 | *MAT1-1* | 1.5 | 10 | HRS09127 | *MAT1-2* | 10 | 1.5 | 325 |
| Pop2 | Pop1_451 | *MAT1-1* | 4 | 7 | Pop1_155 | *MAT1-2* | 3 | 1 | 88 |
| Pop3 | NB29/HRS09122 | *MAT1-2* | 8 | 2 | HRS14051 | *MAT1-1* | 1.5 | 5 | 89 |

Three *Ptt* isolates possessing single virulence QTL for Skiff (Pop2_92) or Prior (Pop2_72 and Pop2_33) were selected from Pop2 based on their genotype within the respective QTL regions and their disease reaction scores. These isolates, along with isolate Pop1_451, one *Ptt* isolate selected from Pop3 (Pop3_55) and isolates NB50 (virulent on Skiff and avirulent on Prior) and NB85 (avirulent on Skiff and virulent on Prior) (Fowler 2018) were used to identify corresponding susceptibility or resistance QTL in barley by phenotyping across recombinant inbred line (RIL) population Prior/Skiff (PS).

### Plant materials

Four barley cultivars, Beecher, Commander, Prior and Skiff, were used to conduct phenotyping of Pop1, Pop2 and Pop3 fungal populations to identify virulence/avirulence QTL in *Ptt*. Beecher and Commander were used as the resistant and susceptible control, respectively.

The Skiff/Prior RIL population (generation F8) consisting of 250 individuals used to identify susceptibility/resistance QTL was developed by Hermitage Research Facility, Department of Primary Industries, Warwick, Queensland, Australia (Fowler 2018).

### Genotyping of pathogen and RIL populations

Progeny and parental fungal isolates were grown on potato dextrose agar (PDA) medium (20 g/L PDA; Biolab Merck, Darmstadt, Germany) at 22 °C for 10 days. Mycelium from each fungal isolates was scraped from the PDA medium and freeze dried for 24 h. Freeze-dried samples were sent to Diversity Arrays Technology Pty. Ltd. (Canberra, ACT, Australia) for DNA extraction and DArTseq™.

Barley leaf tissue was collected from young leaves of F6 plants. DNA was extracted using the Cetyltrimethylammonium bromide (CTAB) method recommended by Diversity Arrays Technology (DArT™) (http://www.diversityarrays.com) for plant tissues (Fowler 2018). The RIL population was genotyped by DArT™ using next generation sequencing platforms to generate marker data from DArTseq™ single nucleotide polymorphisms (SNPs). These markers were used for the genetic map construction of the RIL population.

### Phenotypic Evaluation and Disease Assessment

Seedling stage assessment of all the isolates, except NB50 and NB85, were conducted following the completely randomized block design in a controlled environment room (CER) at the University of Southern Queensland, Australia, with three replicates (Dahanayaka et al. 2022; Dahanayaka&Martin 2023). Barley genotypes were grown in pots with 5 cm diameter and 14 cm height. Each pot contained four plants each of four barley genotypes (Beecher, Commander, Prior and Skiff). Barley genotypes were grown with 12 h day and night cycles at 23 °C and 17 °C respectively, for 14 days at 75 % humidity.

Preparation of conidial suspensions and plant inoculations was as described in Dahanayaka et al. (2022). Fourteen days after planting, 2.5 mL (10000 conidia/mL) of conidial suspension were sprayed on each pot. Parental isolate HRS09127 and *Ptt* isolate HRS10136, with known pathotype (Dahanayaka&Martin 2023), were used as control isolates for each inoculation run to monitor differences across runs. For the PS RIL population, Commander and Beecher were used as the susceptible and resistant controls, respectively. Inoculated pots were incubated in the dark for 24 h at 95 % humidity with a temperature of 23 ± 1 °C. After 24 h, plants were transferred to the same CER mentioned above for nine days. Nine days after inoculation, disease reaction scores on the second leaf was scored using a 1 (highly resistant) to 10 (highly susceptible) scale (Tekauz 1985).

The seedling assays for the NB50 and NB85 isolates were conducted at the Hermitage Research Facility, Department of Primary Industries, Warwick, Queensland, Australia. For the seedling experiments, seeds were sown in 10 cm pots, with five seeds per genotype placed in three designated positions within each pot. A Latin square design was implemented. Plants were grown in a CER at 14 °C night / 24 °C day temperatures with a 12 h photoperiod (Fowler 2018). Inoculations and scoring of plants were conducted as above.

For the adult phenotyping experiments, disease nurseries were established during the winter of 2016 in hill plots at the Hermitage Research Facility, Department of Primary Industries, Warwick, Queensland, Australia, arranged in a randomized complete block design. Separate nurseries were maintained for NB50 and NB85 isolates, ensuring a minimum distance of 500 m to prevent cross-contamination. Hill plots were sown with approximately five seeds per genotype, with designated susceptible spreader genotypes facilitating natural disease dispersal. The inoculation process involved two stages: first, a pre-season disease increase block was inoculated using a mycelial broth spray; second, infected plant material was spread over disease spreaders in the nursery. Disease development was promoted using regular irrigation events. Disease severity in adult plants was recorded on a 1 to 10 scale, following the approach of Saari&Prescott (1975). Data collection was conducted at two points (four and five months after inoculation) for each experiment (Fowler 2018). For all disease score data, the averages of replicates were used for QTL analyses.

### Chi-square analyses

Chi-square (x^2^) tests were performed on phenotypic data from each fungal population and for the RIL population to estimate the number of genes involved in virulence (fungal populations) or resistance (RIL population). For fungal populations, isolates with disease scores less than 6 (<6) were classified as avirulent and those ≥ 6 as virulent. Similarly, RILs with scores <6 were grouped as resistant and those with scores ≥ 6 as susceptible.

### Genetic map construction and QTL analysis

Genetic marker data of the fungal and barley populations generated through DArTseq™ were qualitatively filtered using Microsoft Excel (Dahanayaka et al. 2022) and JoinMap (Ooijen 2018). For the fungal mapping populations, markers non-polymorphic for the parental isolates were removed. Markers and isolates with less than 90% coverage and minor allele frequency less than 1 were removed. Clonal isolates resulting from the nuclear division occurring at the second phase of meiosis of *Ptt* were detected by the *clonecorrect* function in *poppr* package version 2.8.3 (Kamvar et al. 2014) in RStudio version 3.0.2 (R 2013). After quality filtering, DArTseq™ markers from both the fungal and barley populations were grouped into linkage groups using the *make linkage group* function in MapManager QTXb20 version 2.0 (Manly et al. 2001) with a *P* = 0.05 search linkage criterion. Markers were re-ordered using RECORD (Van Os et al. 2005). The final genetic map of individual populations were obtained by manual map curation (Lehmensiek et al. 2009). Marker positions of the resulting genetic maps of the *Ptt* populations were compared with marker positions of the *Ptt* reference genome W1-1 (BioSample SAMEA4560035 available under PRJEB18107 BioProject) to confirm the order of the markers within each linkage group. Short reads (̴ 62 bp) of DArTseq™ marker sequences were mapped to the W1_1 genome using the bowtie2 (Langmead&Salzberg 2012) function in Galaxy (https://usegalaxy.org.au/) and the resulting BAM file with marker positions were compared with the marker order of each linkage group. The linkage groups were then named based on the corresponding chromosome number of the W1_1 reference genome. Marker positions of the PS genetic map were compared with barley reference genome Morex (BioSample SAMEA7384724 available under PRJEB40589 BioProject) and the linkage groups named.

QTL analyses was performed using the composite interval mapping method in Windows QTL Cartographer version 2.5 (Wang 2007). Experiment-wise logarithm of the odds (LOD) threshold values at the 0.05 significance level were calculated with 1000 permutation tests for each trait (Churchill&Doerge 1994; Doerge&Churchill 1996). The phenotypic variances explained by each QTL (R^2^) were calculated. The final QTL figures were drawn using MapChart version 2.32 (Voorrips 2002).

### QTL accumulation

After identifying QTL associated with virulence for Skiff and Prior in Pop1, progeny isolates were grouped based on the QTL they harboured and the number of QTL per isolate. Disease reaction scores were recorded for each group. A Student’s t-test was performed to assess the relationship between QTL accumulation and disease reaction scores. The same analyses were conducted for QTL identified for Skiff susceptibility in SP RIL population.

### RNA extraction and sequencing

Single QTL isolate Pop2_51 possessing the major Chr 5 QTL associated with virulence on Prior and avirulent isolate Pop2_4 from Pop2 were used for RNA sequencing analysis. The second leaves from 14-day old Prior seedlings were cut into 3 cm segments and placed in Petri dishes containing agar, supplemented with 50 mg/L of benzimidazole. Each Petri dish contained three leaf segments. Three droplets of 5 μl conidia solution (1 × 10³ conidia/mL in 0.05% Tween-20) were applied onto each leaf segment in the Petri dishes. Petri dishes were incubated in GEN1000 IN Plant Incubation Chamber (Conviron, Manitoba, Canada) with 12 h day and night cycles at 17°C. Leaf segments were collected at 2, 4 and 7 days post inoculation (dpi) from Petri dishes. All leaf segments were snap-frozen in liquid nitrogen and stored at −80°C until RNA extraction. For each timepoint, three replicates were used in total 18 samples. Leaf samples from Pop2_51 and Pop2_4 were designated as virulent and avirulent, respectively.

RNA was extracted using a miRNeasy Mini Kit (Qiagen, Venlo, The Netherlands) as per the manufacturer’s instructions. RNA samples were quantified using the NanoDrop 2000 mentioned above and 20 μg of each RNA sample was sent to Australian Genome Research Facility (Victoria, Australia) and sequenced using TruSeq stranded mRNA on a NextSeq500 instrument with 100 million read depth. The sequences were paired-end reads with 150 bp each. The sequence read data were deposited in the NCBI under submission number SUB15710218.

### RNA sequencing for barley and Ptt for candidate gene detection

RNA sequencing reads were quality checked with FastQC v 0.12.0 (Andrews 2010) and the adapters were trimmed with Cutadapt v 5.1 (Martin 2011) with the adapter sequence “ACTGTCTCTTATACACATCT”. Forward (F) and reverse (F) RNA sequences for each sample were concatenated to obtain one F and R sequence for each sample and aligned to the barley reference genome MorexV3 (Mascher et al. 2021) and W1-1 Ptt reference genome (Syme et al. 2018) using the RNA-seq aligner Star v 2.7.0 (Dobin et al. 2013) with quantMode set to GeneCounts.

Gene counts resulting per sample were imported to DESeq2 v1.30.1 (Love et al. 2014) in RStudio v 4.5.1 (RCoreTeam 2025). Eighteen samples were grouped into six groups with three replicates each. The dataset was filtered by removing genes for which fewer than two samples within each group had read counts less than 10. Differential expression analysis (DEA) between sample groups was conducted on the normalized data, with adjusted p-values calculated using the Benjamini and Hochberg method to control the false discovery rate (FDR < 0.05) (Benjamini&Hochberg 1995). Principal Component Analysis (PCA) on the expression profiles of all samples was performed using the *prcomp* function in RStudio and the plot was visualised by ggplot2 (Wickham 2011). The candidate virulence genes from the pathogen and the candidate resistant genes from the host were detected manually looking at significantly expressed genes identified from DEA within the respective QTL regions. Candidate genes identified from *Ptt* were analysed using EffectorP to predict their potential as effector proteins (Sperschneider&Dodds 2022).

## Results

### Genotyping of fungal populations

For the fungal populations Pop1, Pop2 and Pop3, a total of 4809, 8775 and 6384 SNPs and SilicoDArT markers, respectively were obtained through DArTseq™. After filtering markers for non-polymorphism and 10% missing values, 1296, 361 and 418 were retained from Pop1, Pop2 and Pop3, respectively. Clonal isolates were detected only in Pop1. Thus, after removing clonal isolates, all the isolates from Pop2 (*n* = 88), Pop3 (*n*= 89) and 305 *Ptt* isolates from Pop1 were retained for genetic map construction.

### Genotyping of the barley population

A total of 5556 markers were obtained for the barley RIL (PS) population. After quality filtering mentioned above 1795 markers were retained for the genetic map construction of RIL population. A total of 250 RILs were used in further analyses.

### Phenotyping of Ptt isolates and RILs

Disease reaction scores of the Pop1 (Figure 2A & B), Pop2 (Figure 2C & D) and Pop3 (Figure 2E & F) on Skiff and Prior ranged from avirulent to virulent with Skewness ranging from - 0.4654 to 0.1124 and transgressive segregation was observed (Table 2). The average disease reaction score for Skiff was 6 in Pop1 and Pop3, and 5 in Pop2. For Prior, the average score across all populations was 5.

**Figure 2.**
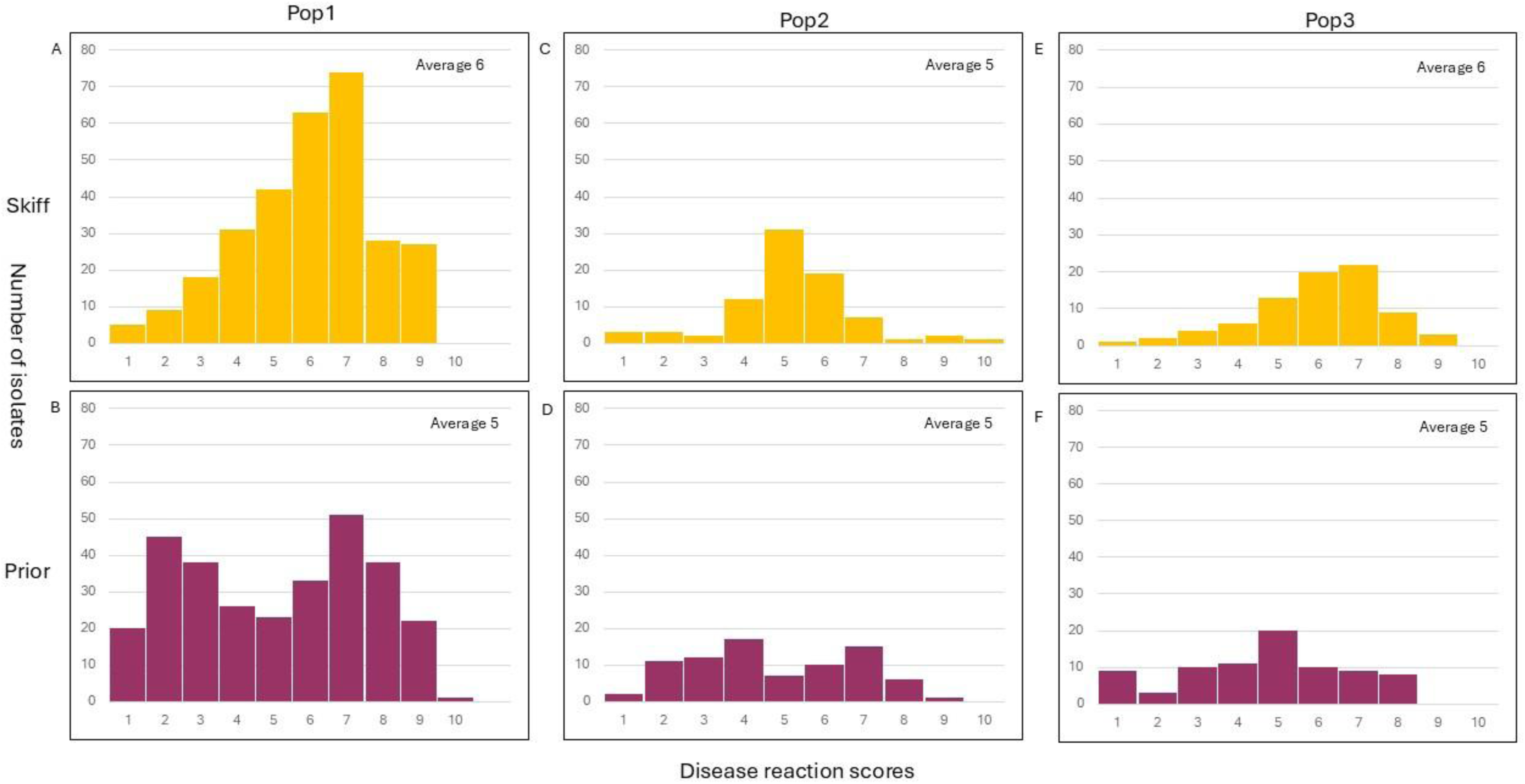
Distribution of disease reaction scores of progeny isolates of Pop1(NB81/HRS09127) on barley varieties (A) Skiff and (B) Prior, Pop2 (Pop1_451/Pop1_155) on Skiff (C) and Prior (D) and Pop3 [(NB29/HRS09122)/HRS14051] on Skiff (E) and Prior (F).

**Table 2.** Disease reaction score ratio of barley varieties Skiff and Prior after inoculation with isolates of the fungal bi-parental mapping populations

|  | Skiff | Prior | 1:1 <sup>a</sup> |  | 3:1 <sup>b</sup> |  | Skewness |  |
| --- | --- | --- | --- | --- | --- | --- | --- | --- |
|  | Vir:Avir <sup>c</sup> | Vir:Avir | Skiff | Prior | Skiff | Prior | Skiff | Prior |
| Pop1 | 192:104 | 145:152 | 0.0001<sup>d</sup> | 0.6846 | 0.0001<sup>d</sup> | 0.0001 | -0.4654 | -0.0611 |
| Pop2 | 30:51 | 32:49 | 0.0589 | 0.0589 | 0.0001 | 0.0001 | -0.1261 | 0.1124 |
| Pop3 | 54:26 | 27:53 | 0.0017 | 0.0037 | 0.1213 | 0.0001<sup>e</sup> | -0.7076 | -0.1869 |

<sup>a</sup> p value for segregation ration of virulent to avirulent isolates for a single gene
<sup>b</sup> p value for segregation ration of virulent to avirulent isolates for two genes
<sup>c</sup> Ratio between the number of virulent isolates to number of avirulent isolates
<sup>d</sup> Number of virulent isolates > number of avirulent isolates
<sup>e</sup> Number of avirulent isolates > number of virulent isolates

The distribution of disease reaction scores among RILs for isolates Pop1_451, Pop2_33, Pop2_72, Pop2_92, Pop3_55, NB50 and NB85 (Figure 3A-I) followed a bimodal distribution. The distributions with isolate Pop1_451 was strongly skewed (−0.8175) toward virulence (susceptibility on barley) whereas with isolate NB50 the distribution was strongly skewed toward avirulence at both the seedling (0.6122) and adult (0.8737) stages (Table 3). The average disease reaction scores ranged from 5 to 7 for the isolates.

**Figure 3.**
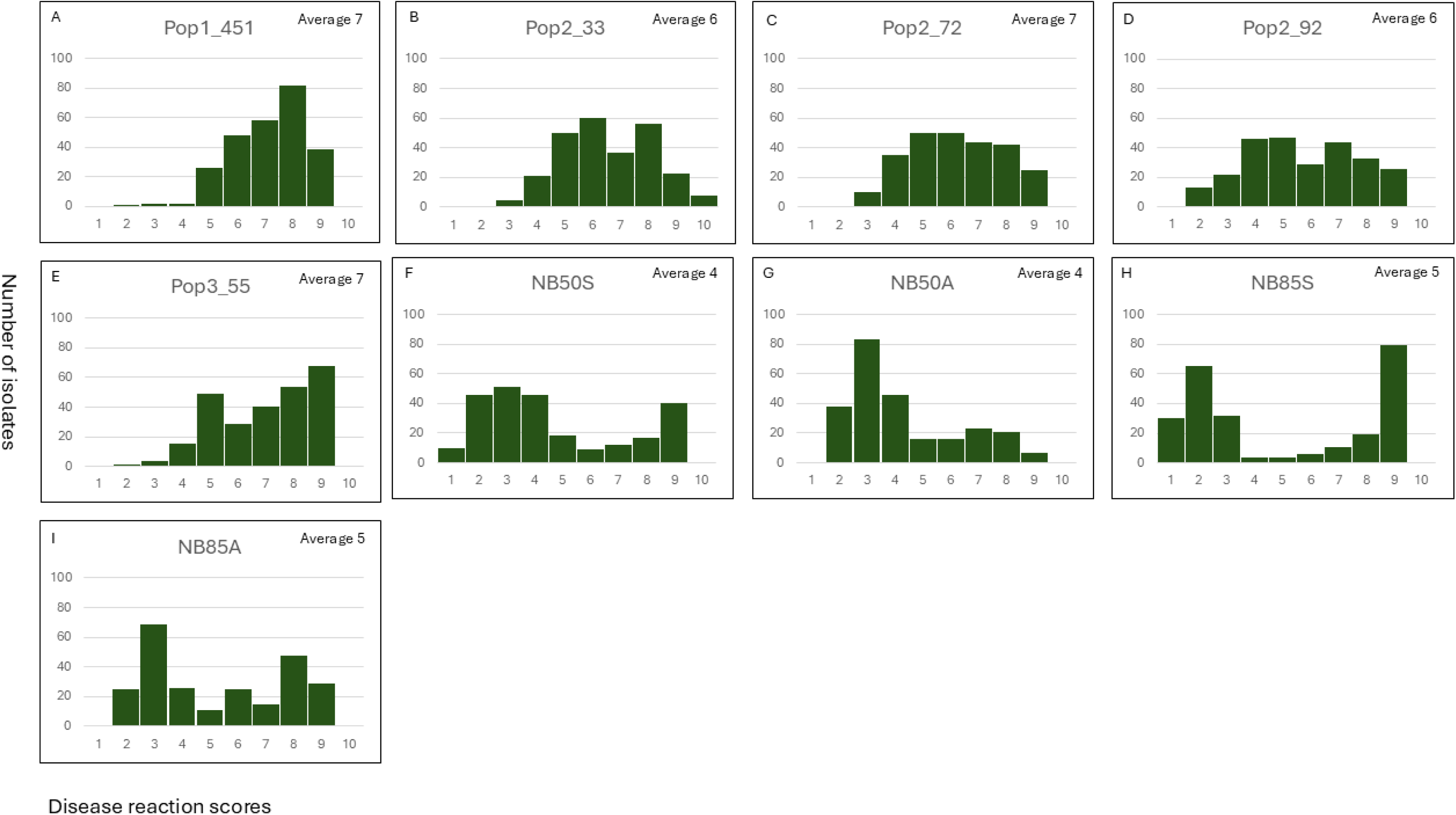
Distribution of disease reaction scores of Skiff/Prior recombinant inbred lines infected with isolates Pop1_451(A), Pop2_33 (B), Pop2_72 (C), Pop2_92 (D), Pop3_55 (E), NB50S (F; seedling), NB50S (G; adult), NB85S (H; seedling) and NB85A (I; adult).

**Table 3.**
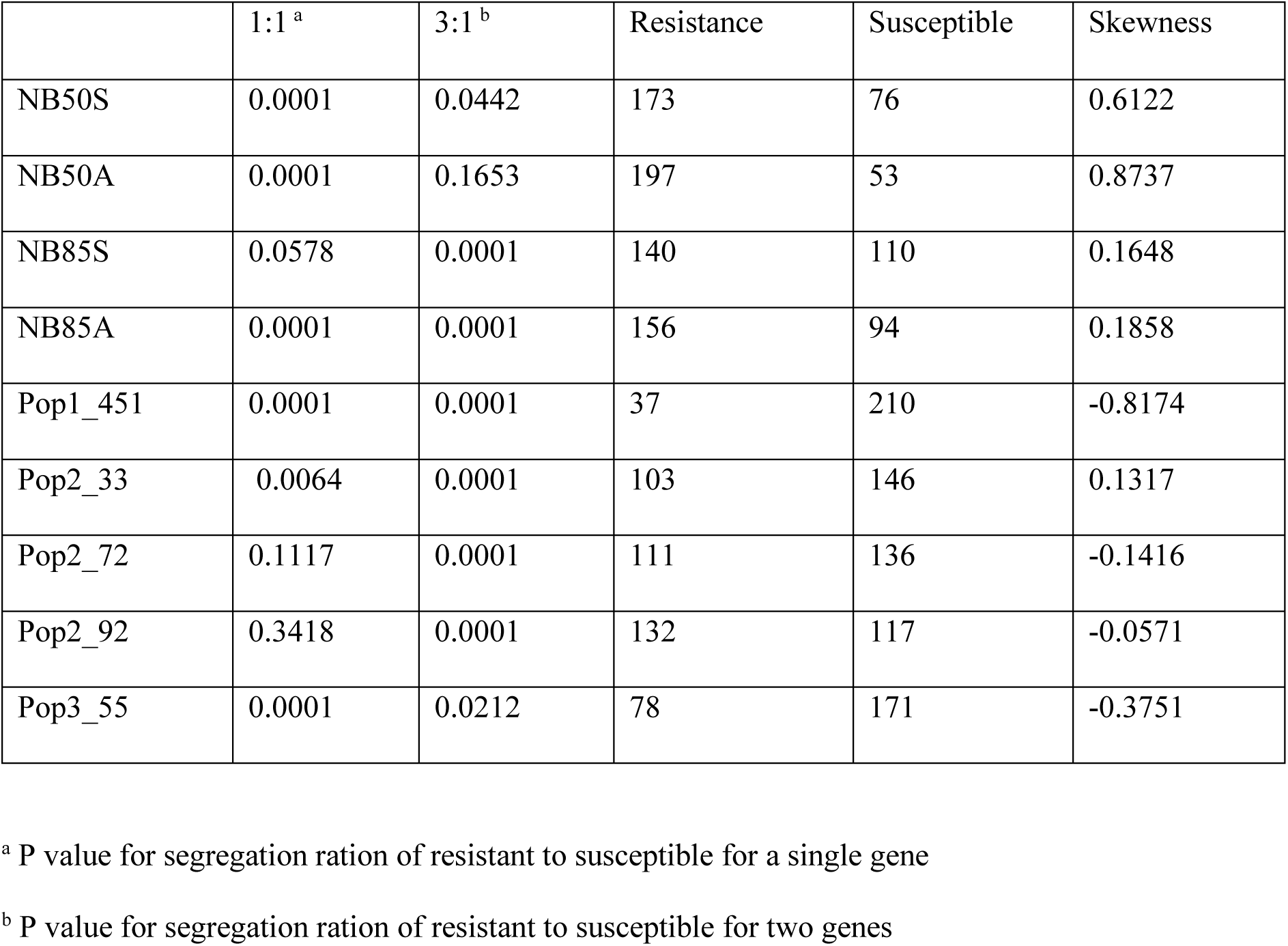
Disease reaction score ratio of recombinant inbred lines of the Skiff/Prior population after infection with different isolates

The segregation of virulent (≥ 6) and avirulent (< 6) isolates based on their phenotypes was evaluated using chi-square (x^2^) tests, expecting 1:1 (single gene) and 3:1 (two genes) ratios for Pop1, Pop2, and Pop3 on both Skiff and Prior (Table 2). For Pop1, segregation on Skiff significantly deviated from both 1:1 and 3:1 expectations (p = 0.001), while no significant deviation was observed for Prior under the 1:1 model. For Pop2, no significant deviation from the expected 1:1 ratio was observed for either Skiff or Prior. For Pop3, segregations on Skiff did not significantly differ from the 3:1 ratio, whereas a significant difference was observed on Prior.

Chi-square analysis of the segregation ratios between susceptible (≥ 6) and resistant (< 6) phenotypic groups for the PS RILs for 1:1 (single genes) model showed no significant difference for NB85 (seedling assay), Pop2_33, Pop2_72 and Pop2_92. For 3:1 (two genes) model no significant difference was observed for both seedling and adult plant assays of NB50 and Pop3_55.

### Genetic map and QTL mapping

The genetic map of Pop1 consisted of 12 linkage groups with 570 non-redundant markers. The genetic maps for Pop2 and Pop3 consisted of 13 and 14 linkage groups with 160 and 217 non-redundant markers respectively. The average distance between flanking markers of Pop1, Pop2 and Pop3 were 3.5, 7.7 and 3.8 cM, respectively (Table 4).

**Table 4.** Information of genetic linkage maps of the fungal bi-parental mapping populations.

| Cross ID | Number of isolates/RILs | Number of non-redundant markers | Size of the genetic map (cM) | Average distance between flanking markers |
| --- | --- | --- | --- | --- |
| Pop1 | 305 | 570 | 2019.6 | 3.5 |
| Pop2 | 88 | 160 | 1229.5 | 7.7 |
| Pop3 | 89 | 217 | 827.7 | 3.8 |
| SP RIL | 250 | 1628 | 1357.9 | 0.8 |

The genetic map of PS RILs consisted of seven linkage groups and 1628 non-redundant markers with 0.8 cM average distance between flanking markers.

The QTL analysis of Pop1 detected three QTL for Prior virulence on Chr 1, 4 and 5 with LOD scores of 6, 11 and 40, respectively (Table 5). The phenotypic variations explained (R^2^) by QTL on Chr 1, 4 and 5 were 8.3, 16.8 and 48%, respectively. For virulence on Skiff, two QTL in Chr 3 and 10 were detected in Pop1 with LOD scores of 22 and 7, respectively. The phenotypic variation explained by the QTL on Chr 3 and 10 was 21.2 and 7%, respectively. The parent contributing towards the increase of the disease reaction scores for Prior was NB81 and for Skiff was HRS09127.

**Table 5.**
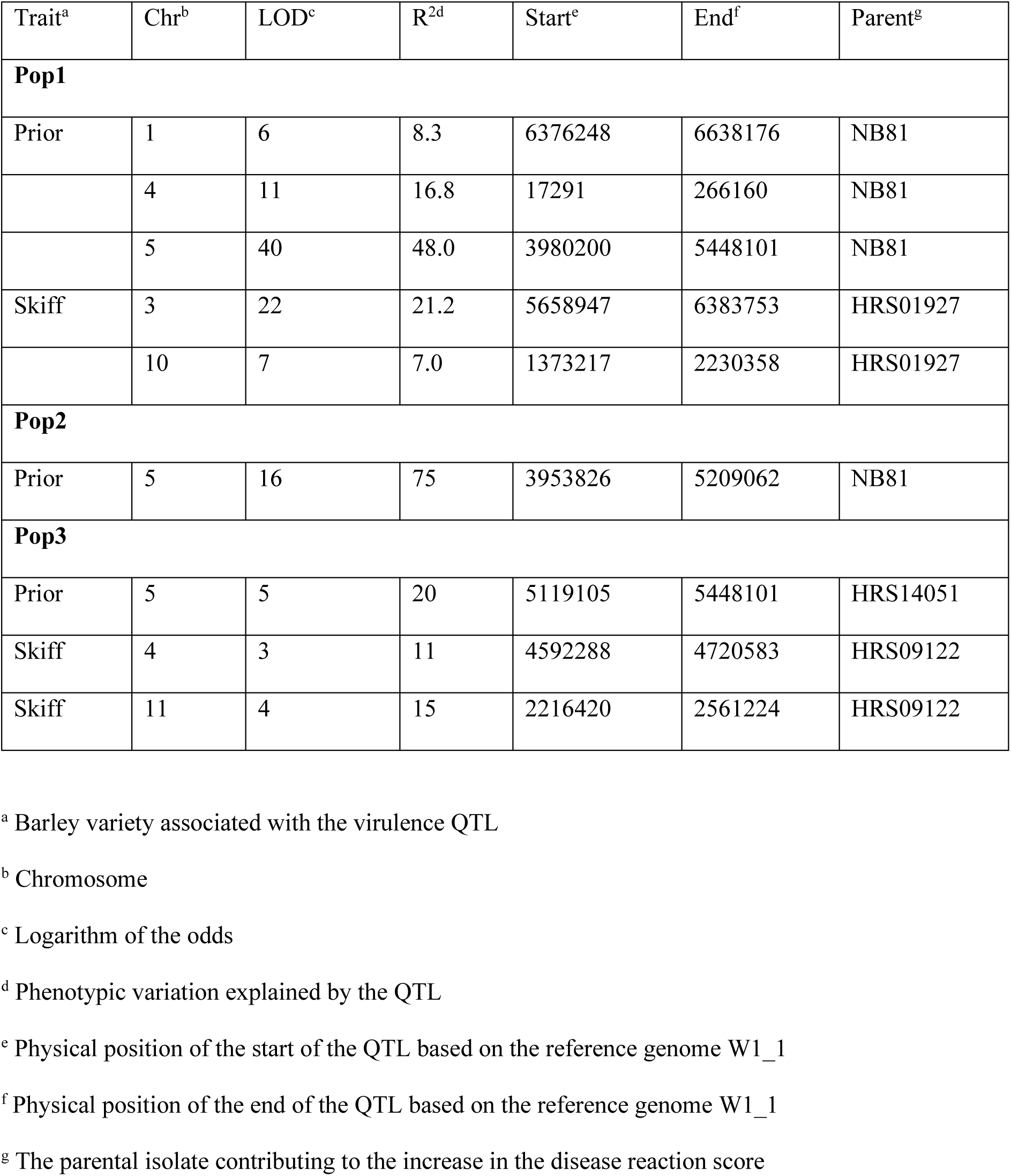
QTL information of the fungal bi-parental mapping populations.

| Trait<sup>a</sup> | Chr<sup>b</sup> | LOD<sup>c</sup> | R<sup>2d</sup> | Start<sup>e</sup> | End<sup>f</sup> | Parent<sup>g</sup> |
| --- | --- | --- | --- | --- | --- | --- |
| **Pop1** |  |  |  |  |  |  |
| Prior | 1 | 6 | 8.3 | 6376248 | 6638176 | NB81 |
|  | 4 | 11 | 16.8 | 17291 | 266160 | NB81 |
|  | 5 | 40 | 48.0 | 3980200 | 5448101 | NB81 |
| Skiff | 3 | 22 | 21.2 | 5658947 | 6383753 | HRS01927 |
|  | 10 | 7 | 7.0 | 1373217 | 2230358 | HRS01927 |
| **Pop2** |  |  |  |  |  |  |
| Prior | 5 | 16 | 75 | 3953826 | 5209062 | NB81 |
| **Pop3** |  |  |  |  |  |  |
| Prior | 5 | 5 | 20 | 5119105 | 5448101 | HRS14051 |
| Skiff | 4 | 3 | 11 | 4592288 | 4720583 | HRS09122 |
| Skiff | 11 | 4 | 15 | 2216420 | 2561224 | HRS09122 |

<sup>a</sup> Barley variety associated with the virulence QTL
<sup>b</sup> Chromosome
<sup>c</sup> Logarithm of the odds
<sup>d</sup> Phenotypic variation explained by the QTL
<sup>e</sup> Physical position of the start of the QTL based on the reference genome W1\_1
<sup>f</sup> Physical position of the end of the QTL based on the reference genome W1\_1
<sup>g</sup> The parental isolate contributing to the increase in the disease reaction score

QTL analysis of Pop2 detected one QTL for Prior virulence on Chr 5 with LOD of 16 and 75% of phenotypic variance explained. No QTL was detected for virulence on Skiff for Pop2. For Pop3, one QTL was detected for virulence on Prior on Chr 5 with LOD of 5 and 20% of the phenotypic variance explained and two QTL for Skiff virulence on Chr 4 and 11 with LOD of 3 and 4 and 11 and 15% of the phenotypic variance explained, respectively.

QTL analyses of the PS RIL population using seven isolates, NB50 (NB50A: adult and NB50S: seedling plant assays), NB85 (NB85A: adult and NB85S: seedling plant assays), Pop1_451, Pop2_33, Pop2_72, Pop2_92 and Pop3_55 detected two major QTL one on Chr 3H and the other on 6H (Figure 4). The QTL *UniSQRpt3Ha,* on Chr 3H of barley was detected after inoculation with Skiff virulent isolates NB50 (seedling and adult stages), Pop2_92 and Pop3_55 with LOD scores ranging from 34 to 62 and phenotypic variance explained ranging from 37 to 56% (Table 6, Figure 4).

**Figure 4.**
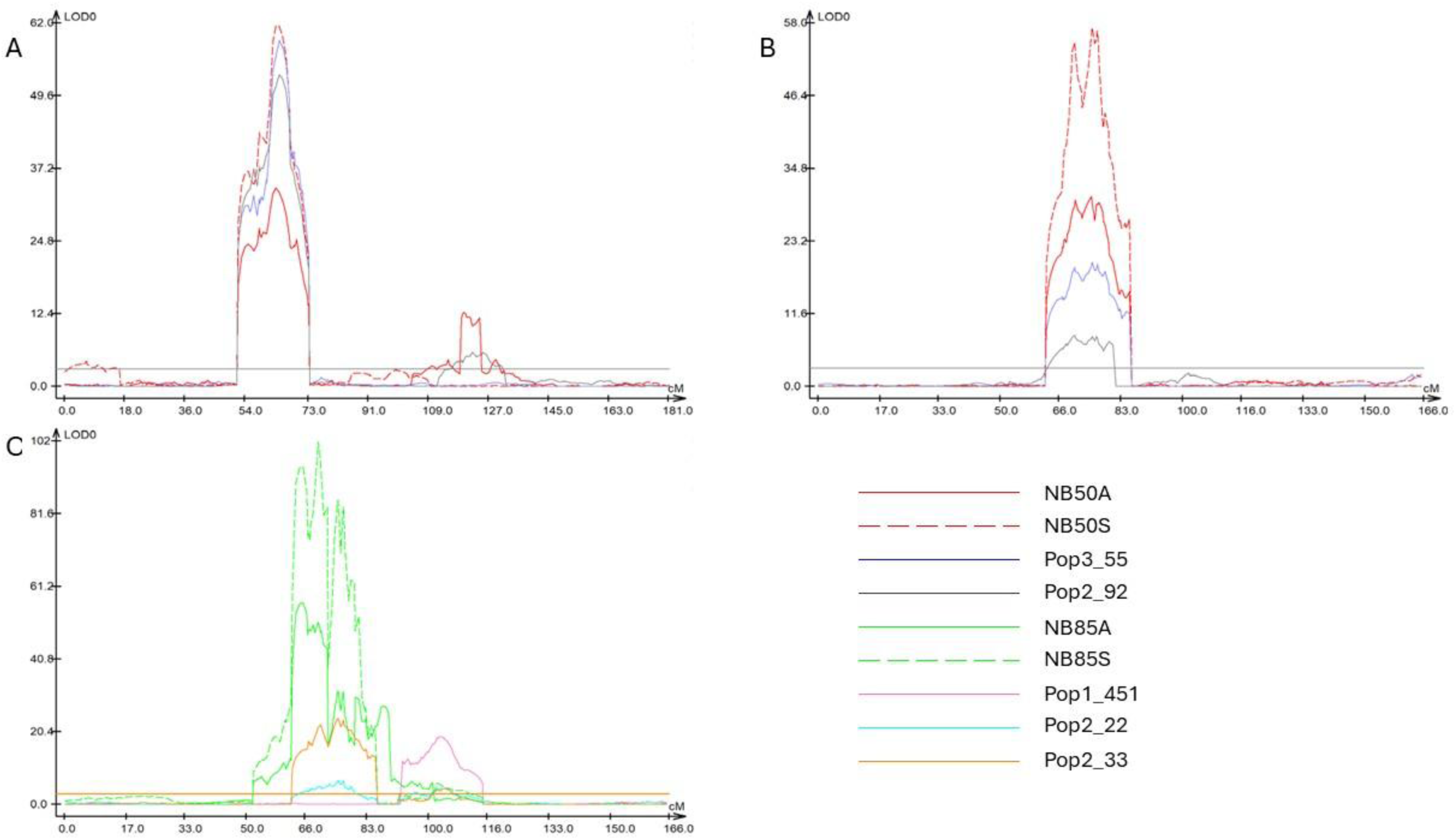
Barley quantitative trait loci (QTL) (p = 0.05) associated with resistance to *Pyrenophora teres* f. *teres*. QTL linked to virulence on Skiff were located on chromosome 3H (A) and 6H (B), and those associated with virulence on Prior mapped to chromosome 6H (C).

**Table 6.** QTL information of the Skiff/Prior recombinant inbred line mapping population.

| QTL<sup>a</sup> | Chr<sup>b</sup> | LOD<sup>c</sup> | R<sup>2d</sup> | Isolate | Stage | Physical map based on the reference genome (bp) |  | R source<sup>e</sup> |
| --- | --- | --- | --- | --- | --- | --- | --- | --- |
|  |  |  |  |  |  | Start | End |  |
| *UniSQRpt3Ha* | 3H | 62 | 37 | NB50 | Adult | 63638040 | 434379440 | Prior |
|  |  | 34 | 28 | NB50 | Seedling |  |  |  |
|  |  | 54 | 55 | Pop2_92 | Seedling |  |  |  |
|  |  | 60 | 56 | Pop3_55 | Seedling |  |  |  |
| *UniSQRpt3Hb* |  | 13 | 14 | NB50 | Adult | 486616920 | 511570680 | Prior |
|  |  | 10 | 3 | NB85 | Adult |  |  | Prior |
|  |  | 6 | 6 | Pop1_451 | Seedling |  |  | Prior |
|  |  | 6 | 5 | Pop2_33 | Seedling |  |  | Prior |
|  |  | 6 | 15 | Pop2_92 | Seedling |  |  | Prior |
| *UniSQRpt6Ha* | 6H | 31 | 23 | NB50 | Adult | 39476640 | 418904240 | Prior |
|  |  | 58 | 33 | NB50 | Seedling |  |  | Prior |
|  |  | 20 | 13 | Pop3_55 | Seedling |  |  | Prior |
| *UniSQRpt6Hb* |  | 102 | 59 | NB85 | Seedling | 21946760 | 400093440 | Skiff |
|  |  | 57 | 35 | NB85 | Adult |  |  | Skiff |
|  |  | 25 | 24 | Pop2_33 | Seedling |  |  | Skiff |
|  |  | 7 | 9 | Pop2_72 | Seedling |  |  | Skiff |
|  |  | 9 | 5 | Pop2_92 | Seedling | 39476640 | 418904240 | Skiff |
| *UniSQRpt6Hc* |  | 19 | 26 | Pop1_451 | Seedling | 436070800 | 503364360 | Skiff |
|  |  | 5 | 9 | Pop2_33 | Seedling |  |  | Skiff |
|  |  | 5 | 9 | Pop2_72 | Seedling |  |  | Skiff |

<sup>a</sup> Quantitative trait loci (QTL)
<sup>b</sup> Chromosome
<sup>c</sup> Logarithm of the odds
<sup>d</sup> Phenotypic variation explained by the QTL
<sup>e</sup> Parental isolate contributing the resistance

A minor QTL, *UniSQRpt3Hb* was also detected on 3H after inoculation with NB50A, NB85A, Pop1_451, Pop2_33, Pop2_92. For these two QTL on Chr 3H, the disease reaction scores were higher in individuals having the Skiff alleles compared to those having Prior alleles, hence the resistance source was identified as Prior.

The major QTL in Chr 6H had two peaks within the QTL region, hence they were designated as *UniSQRpt6Ha* and *UniSQRpt6Hb*. The QTL *UniSQRpt6Ha* was detected after infection with NB50A, NB50Sand Pop3_55 (Figure 4). The LOD values of the QTL ranged from 20 to 58 with 13 to 33% phenotypic variation explained. The resistance source for the QTL was Prior.

The QTL *USQRpt6Hb* was detected after infection with NB85a, NB85s, Pop2_33, Pop2_72 and with LOD values ranging from 7 to 102 and phenotypic variation accounting for 5 to 59%.

The QTL *UniSQRpt6Hc*, identified after infection with Pop1_451, and Pop2_72 with LOD value between 5 and 19 and 9 to 26% of the phenotypic variation explained. The resistance source for the QTL was Skiff.

### QTL accumulation

The QTL accumulation effects for the QTL on Chr 3 and 10 associated with virulence on Skiff (Figure 5A) and QTL on Chr 1, 4, and 5 associated with virulence on Prior were examined (Figure 5B). Progeny isolates harbouring either QTL on chr 3 or 10 were significantly different (p = 0.001) from isolates that do not harbor any QTL (Table 7). When both QTL were present, the average disease reaction score was 7, and significantly different (p = 0.01) from harbouring either of the single QTL. The average disease reaction score of progeny isolates harbouring QTL on Chr 5, either as a single QTL or coupled with others, was significantly higher (p = 0.001 or 0.01) than isolates possessing QTL on Chr 1 and 4 (Table 8). The highest disease reaction score was observed when all three QTL were present.

**Figure 5.**
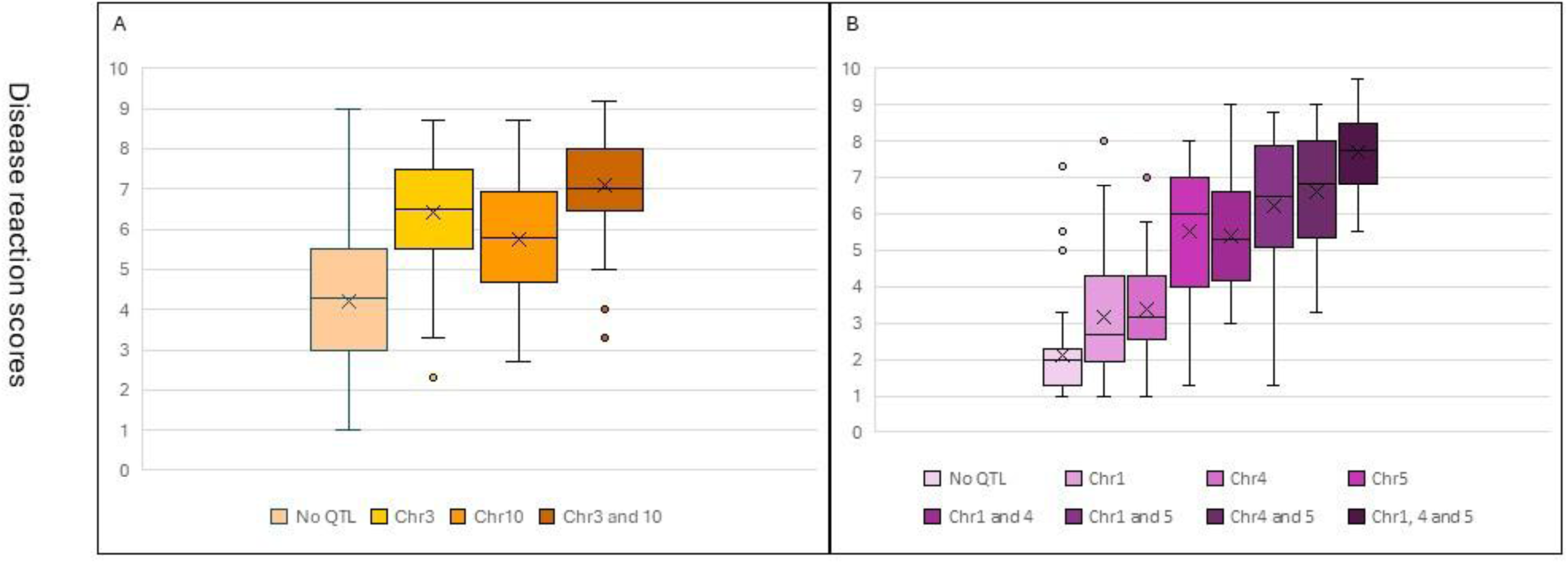
Accumulation effects of virulence QTL of the *Pyrenophora teres* f. *teres* progeny isolates on barley cultivars Skiff (A) and Prior (B).

**Table 7.** Average disease reaction scores and their significance during QTL accumulation for Skiff virulence in fungal population Pop1.

|  | No QTL | Chr3 | Chr10 | Average |
| --- | --- | --- | --- | --- |
| No QTL |  |  |  | 4.2 |
| Chr3 | 3.1E-14 |  |  | 6.4 |
| Chr10 | 6.7E-08 | 0.0099 |  | 5.7 |
| Chr3 and 10 | 5.2E-22 | 0.0038 | 8.9E-08 | 7.1 |

Chr – chromosome

**Table 8.** Average disease reaction scores and their significance during QTL accumulation for Prior virulence in fungal population NB81/HRS09172.

|  | No QTL | Chr1 | Chr4 | Chr5 | Chr1 and<br>4 | Chr1<br>and 5 | Chr4<br>and 5 | Average |
| --- | --- | --- | --- | --- | --- | --- | --- | --- |
| No<br>QTL |  |  |  |  |  |  |  | 2.1 |
| Chr1 | 0.0042 |  |  |  |  |  |  | 3.1 |
| Chr4 | 7.4E-05 | 0.6054<sup>ns</sup> |  |  |  |  |  | 3.4 |
| Chr5 | 3.8E-15 | 3.0E-06 | 2.6E-06 |  |  |  |  | 5.5 |
| Chr1<br>and 4 | 1.5E-14 | 1.2E-05 | 3.7E-06 | 0.8692<sup>ns</sup> |  |  |  | 5.4 |
| Chr1<br>and 5 | 3.7E-22 | 1.1E-10 | 1.5E-10 | 0.0800<sup>ns</sup> | 0.00725 |  |  | 6.2 |
| Chr4<br>and 5 | 1.0E-20 | 4.2E-10 | 7.3E-11 | 0.0174 | 0.0126 | 0.3542<sup>ns</sup> |  | 6.6 |
| Chr1, 4<br>and 5 | 6.7E-36 | 1.3E-19 | 1.6E-22 | 3.5E-08 | 5.7E-09 | 3.9E-05 | 0.003 | 7.7 |

Chr Chromosome
<sup>ns</sup> not significant

PS RILs harbouring susceptible QTL on either Chr 3H, 6H or both showed significant increase in disease reaction scores (p = 0.001 or 0.01) compared to those that did not harbour any QTL, with the exception of RILs inoculated with Pop2_92 (Figure 6). The 6H QTL for the Pop2_92 inoculated RILs was a minor QTL and lines having this QTL did not differ from those without any susceptible QTL. For inoculations with the NB50 isolate, there was no difference in the disease reaction score for RILs having either the 3H or 6H QTL at both seedling and adult plant stages.

**Figure 6.**
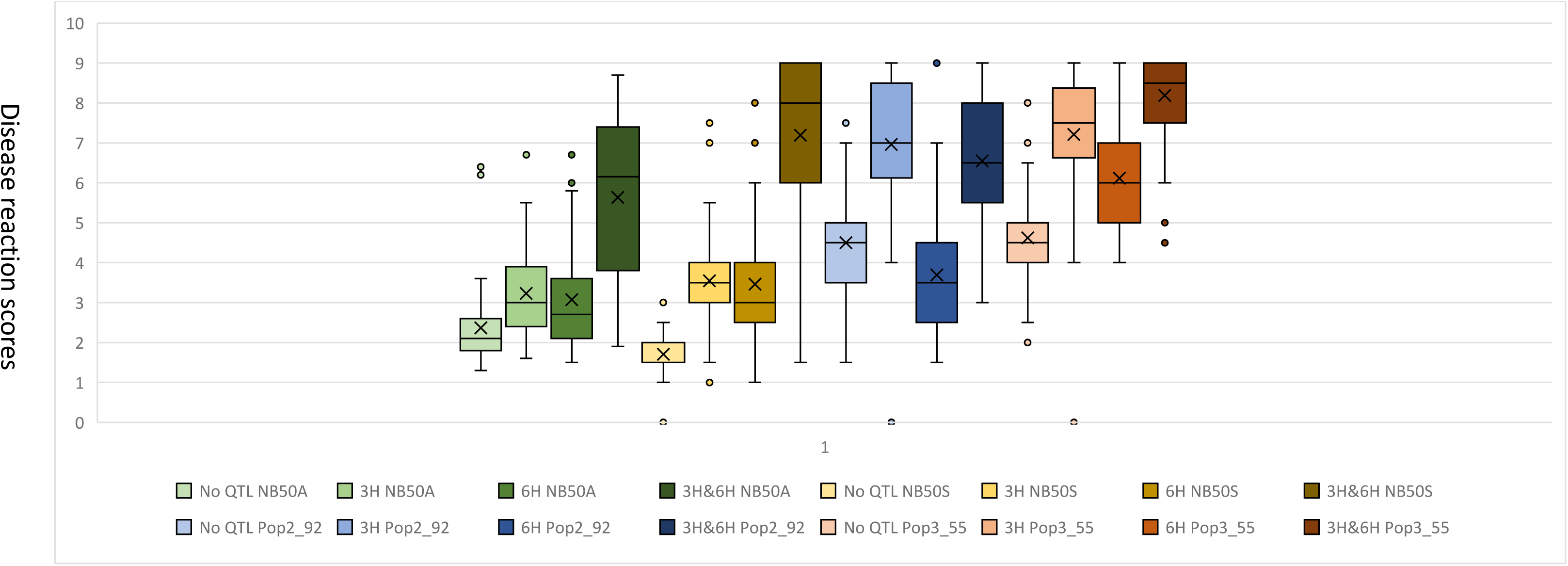
Accumulation effects of susceptible QTL *UniSQRpt3Ha* (3H) and *UniSQRpt6Ha* (6H) in the Prior/Skiff recombinant inbred line population for *Pyrenophora teres* f*. teres* isolates possessing QTL associated with virulence on the Skiff barley cultivar.

### Candidate virulence gene detection of Ptt

*In-planta* transcriptomic analysis was conducted using virulent (Pop2_51) and avirulent (Pop2_4) isolates of Pop2. The virulent isolate contained the major QTL in Chr 5 for virulence on Prior while this QTL was absent in the avirulent isolate. PCA of RNA-seq data from the virulent and avirulent *Ptt* isolates across different post-inoculation time points revealed distinct clusters for D2 samples and another cluster for D4 and D7 samples (Figure 7). PC1, which explained the largest proportion of variance (∼16.2%), was significantly influenced by time post-inoculation (ANOVA, p = 0.001). PC2, accounting for ∼14.5% of the variance was significantly associated with the virulence of the *Ptt* isolates (ANOVA, p = 0.01).

**Figure 7.**
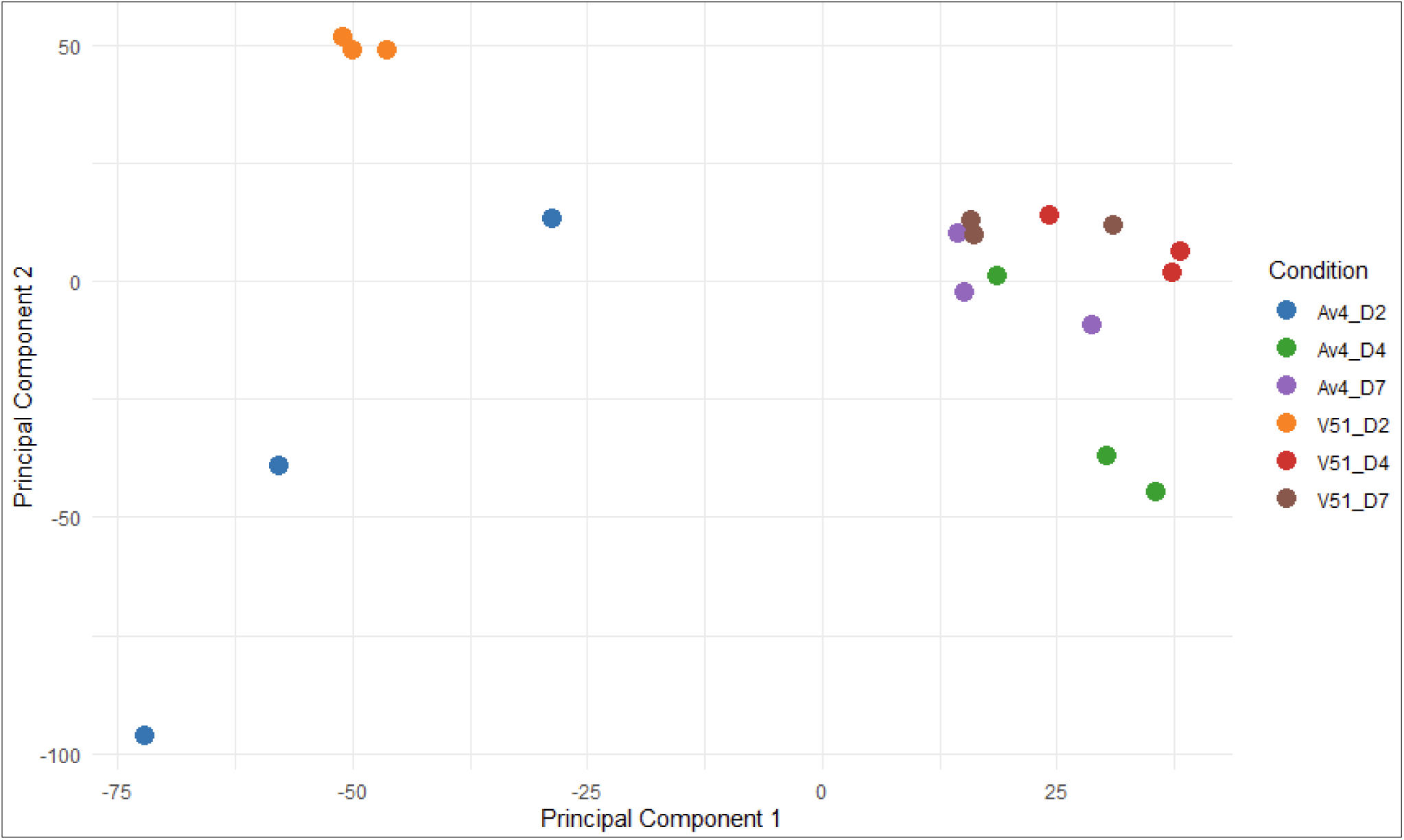
Principal Component Analysis (PCA) of RNA-seq data from virulent (Pop2_51; V51) and avirulent (Pop2_4; Av4) *Pyrenophora teres* f*. teres* isolates across post-inoculation time points.

At 2D, 4D, and 7D post-inoculation; 827, 46, and 28 genes were significantly differentially expressed in the virulent isolate compared to the avirulent isolate, respectively (Supplementary Table 1-3). Five candidate genes, which were significantly differentially expressed in the virulent isolate compared to the avirulent, were located within the Chr 5 QTL. Four of the five genes, PTTW11_06604, PTTW11_06608, PTTW11_06615 and PTTW11_06666 were identified at D2 while PTTW11_06728 was differently expressed at D4 and D7. Among these, PTTW11_06608 and PTTW11_06728 were predicted to encode cytoplasmic and apoplastic effector proteins, respectively.

### Candidate resistance gene detection of barley

PCA of RNA-seq data from both susceptible and resistant reactions to Prior across multiple post-inoculation time points revealed distinct clusters. These clusters clearly separated samples by time points as well as host response (susceptible/resistant) (Figure 8). PC1, which accounted for the largest proportion of variance (∼37.9%), was significantly influenced by both time post-inoculation (ANOVA, p = 0.001) and host response (susceptible/resistant) (ANOVA, p = 0.01). PC2 explained 15.7% of the variance and was predominantly driven by genotype (ANOVA, p = 0.01).

**Figure 8.**
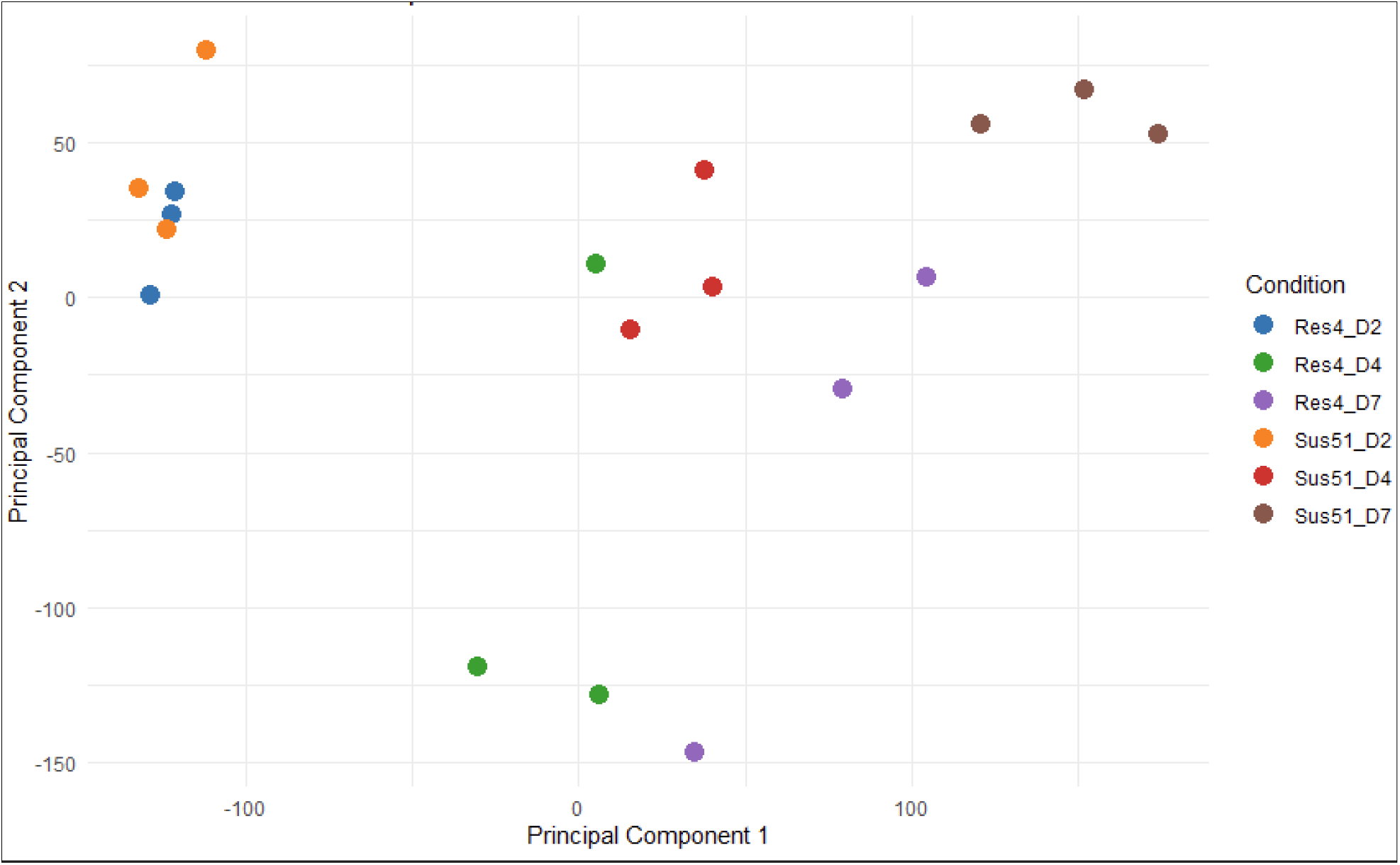
Principal Component Analysis (PCA) of RNA-seq data from the barley cultivar Prior during susceptibility (Sus51) and resistance (Res4) reactions caused by inoculation with virulent (Pop2_51; Sus51) and avirulent (Pop2_4; Res4) *Pyrenophora teres* f. *teres* isolates across post-inoculation time points.

Transcriptomic profiles of susceptible compared to resistant reactions on Prior at D2, 4D, and 7D post-inoculation detected 549, 481, and 4844 significantly differentially expressed genes, respectively (Supplementary Table 4-6). For QTL, *UniSQRpt3Ha* (n = 233), *UniSQRpt3Hb* (n = 34) and *UniSQRpt6Ha* (n = 313) 580 significantly differentially expressed genes in resistant compared to the susceptible reactions were identified for Prior (Table 9).

**Table 9.** Number of candidate genes corresponding to the three barley QTL detected in this study identified through RNA sequencing of samples collected at two (D2), four (D4) and seven (D7) days after inoculation

| QTL | Chr | D2 | D4 | D7 | Total |
| --- | --- | --- | --- | --- | --- |
| *UniSQRpt3Ha* | 3 | 24 | 27 | 182 | 233 |
| *UniSQRpt3Hb* | 3 | 3 | 4 | 27 | 34 |
| *UniSQRpt6Ha* | 6 | 30 | 22 | 261 | 313 |
| *Total* |  | 57 | 53 | 470 | 580 |

QTL quantitative trait loci
Chr Chromosome

## Discussion

The interaction between plant pathogens and their hosts is often described as an “arms race”. This dynamic involves continuous adaptations and counter-adaptations by both the pathogen and the plant. Pathogens evolve new strategies to infect and overcome plant defences, while plants develop new resistance mechanisms to fend off these attacks. Previous studies have identified genomic regions associated with the barley-*P. teres* f*. teres* pathosystem by separately identifying genomic regions associated with either virulence/avirulence or susceptibility/resistance. In the current study multiple genomic approaches were integrated to identify genomic regions associated with virulence of *Ptt* and linking these to corresponding genomic regions associated with susceptibility in the barley genome. The findings of this study advance our understanding of the genes involved in the interaction between barley and *P. teres*. The pathogen, *P. teres*, has been less extensively studied compared to the host (barley) due to the labour-intensive nature of developing crosses, collecting ascospores, and phenotyping individual isolates. In this study, we employed the largest *P. teres* mapping population to date, comprising of 305 isolates, along with two additional populations of 88 (Pop2) and 89 (Pop3) isolates, to identify QTL in the pathogen. Selected single QTL isolates from Pop2, having only one virulence QTL, confirm that the susceptibility QTL identified in the host correspond to the respective virulence genes in the pathogen.

The QTL mapping of the primary *Ptt* population (Pop1) detected two and three QTL associated with Skiff and Prior, respectively. The QTL on Chr 3 for virulence on Skiff and the QTL in Chr 5 for virulence on Prior were major QTL with phenotypic variation explained of 24 and 40%, respectively. The QTL were found to be co-located with previously identified QTL detected by bi-parental, GWAS and nested-association mapping studies (Martin et al. 2020a; Dahanayaka&Martin 2023), some of which were also associated with virulence on other barley cultivars such as Beecher. These studies and ours suggest that Skiff, Beecher and Prior virulence is located within the same genomic region (Martin et al. 2020a).

Chi-square analyses of the segregation ratios indicated that susceptibility of the PS population to isolate NB50 during both adult and seedling stages, as well as isolate Pop3_55, was mainly due to the interaction of two genes, corresponding to the QTL located on Chr 3H and 6H. In contrast, susceptibility to Pop2_92 was associated with a single QTL on Chr 3H. The chi-square test showed no significant deviation from a 1:1 segregation ratio, indicating that a single gene is likely responsible for this interaction. Even though the QTL associated with NB50 and Pop3_55 susceptibility in SP population were similar, susceptibility-to-resistance ratio between the two isolates were contrasting. Susceptibility-to-resistance ratio for NB50 was 3:1 while for Pop3_55 1:3. This observation indicates that the same QTL in the host might be interacting differently with the pathogen and there could be additional genetic factors influencing the susceptibility/resistance. It could be due to an epistatic interaction where QTL may interact with loci in a way that alters the resistance response to different isolates.

Isolate Pop2_92 derived from the cross between two isolates of Pop1 (451 and 155) inherited the Skiff virulence gene from isolate HRS09127, while isolate Pop3_55 inherited the Skiff virulence from HRS09122. QTL mapping of Pop2 did not detect any QTL for Skiff virulence, however, the original NB81/HRS09127 population detected QTL for Skiff virulence on Chr 3 and 10 while two QTL for Skiff virulence were detected with Pop 3 on Chr 4 and 11. Even though the Skiff virulence QTL were present in different genomic locations, the corresponding QTL in PS was the same (*UniSQRpt3Ha*) for both isolates. Although the host QTL are located in the same genomic region, further work needs to be undertaken to determine whether the same gene or different closely-linked susceptibility genes are interacting with the virulence genes of the two isolates involved.

The chi square analysis of disease reaction score of Pop1_451 and NB85A on the PS RIL population suggested that the host susceptibility is controlled by multiple genes and the distribution was skewed toward susceptibility. This could be due to Pop1_451 and NB85A potentially producing multiple effectors, allowing them to overcome more than one host resistance gene. Furthermore, Pop1_451 possesses the Prior virulence QTL identified on Chr 5 of Pop1. The QTL ranges from 3980200 to 5448101 bp (with respect to W1_1 reference genome), covering a broad range, hence, the isolate may have effector gene clusters, meaning several virulence genes are physically linked to overcome the host resistance.

QTL analyses for susceptibility of PS due to Pop2_33 and Pop2_72, suggest that the susceptibility was due to a single gene interaction on 6H, *USQRpt6Hb*. The chi square analyses of the resistance to susceptibility genotype for Pop2_33 and Pop2_72, also suggest that susceptibility was due to a single gene interaction, as the p value did not deviate for the 1:1 ratio. The isolates Pop2_33 and Pop2_72 were single QTL isolates possessing one QTL for the Prior virulence QTL on Chr 5, unlike its virulent parent Pop1_451 where it may possess an effector gene cluster. Therefore, it can be suggested that the interaction between these isolates with the PS barley population was due to QTL in Chr 5 of the Ptt genome with *USQRpt6Hb* QTL in the PS barley genome.

The QTL, *USQRpt6Ha* and *b* are major QTL co-located in 6H possessing two susceptible sources. This suggests the presence of two closely linked QTL or two alleles for susceptibility in Prior and Skiff to *Ptt*. The susceptibility source for NB50 and Pop3_55 was found to be Skiff while for NB85, pop2_33 and Pop2_72, the susceptibility source was found to be Prior. Similar observations were reported for a barley population developed by Kombar and Rika where Rika was resistant to Ptt isolates 15A but susceptible to 6A, while Kombar showed resistance to 6A and susceptible to 15A (Abu Qamar et al. 2008; Shjerve et al. 2014). These studies reported two QTL for Kombar (*rpt5.k*) and Rika (*rpt5.r*) susceptibility on 6H in repulsion phase. The results of the current study suggest that susceptibility is controlled by more than one separate gene/allele and possibly corresponding to different effectors (Richards et al., 2016).

The QTL region *USQRpt6Ha* and *b* is co-located with the previously reported complex 6H locus *Rpt5/Spt1* (Abu Qamar et al. 2008). The *Rpt5/Spt1* locus in the barley 6H chromosome is reported to present dominant resistance in barley genotypes like CI5791 and dominant susceptibility (Manninen et al., 2006) in genotypes like Rika or Kombar (Abu Qamar et al., 2008 Richards et al., 2016), depending on the specific allele–effector interaction providing resistance or susceptibility based on the combination of alleles and pathogen effectors present (Richards et al., 2016). The region is reported to be involved in both gene-for-gene and inverse gene-for-gene models. A recent study revealed that the necrotrophic effector PttNE1 from Ptt, interacted with a gene designated *SPN1* located at the same 6H chromosome region as the *Rpt5/Spt1* locus, which accounted for 31% of the disease variation following the inverse gene-for-gene model. The current study and above-mentioned studies have revealed the presence of resistance gene/s in the *Rpt5/Spt1* locus which trigger immunity when matched with an avirulence or virulence gene in the pathogen following the gene-for-gene model.

The results of the QTL analyses with host and pathogen phenotypic data from this study provide evidence that both the virulence and susceptibility exhibit additive patterns. In Pop1, where multiple QTL were detected for Skiff and Prior, when *Ptt* isolates harboured more than one gene, the average disease reaction scores were significantly higher than compared to isolates possessing one or two genes. A similar pattern was observed for the PS population for Skiff, where two susceptible loci were detected. The presence of loci on both 3H and 6H resulted in greater resistance than when only one locus was present. This cumulative effect of virulence or susceptibility QTL highlights the importance of QTL combinations in shaping host-pathogen dynamics, consistent with the gene-for-gene concept, where the battle between pathogen virulence/avirulence factors and host resistance/susceptibility genes determines the disease outcome.

*In-planta* transcriptomic analyses identified five candidate genes for the Prior virulence in Ptt. Out of these five, two proteins were uncharacterised proteins (PTTW11_06608 and PTTW11_06728). The remaining proteins were PTTW11_06604; TPMT multi-domain protein; PTTW11_06615; MOSC domain containing protein and PTTW11_06666; Rhodopsin domain-containing protein. Future genome editing studies will be conducted to confirm the gene/s linked to the Prior virulence in Ptt. The major QTL regions associated with Prior resistance are around 370 Mbp, containing more than 500 candidate genes. Future fine mapping using whole-genome sequencing of individuals from the RIL population will be conducted to further narrow down candidate regions.

In conclusion, this study represents a significant advancement in our understanding of the genetic interactions within the barley-*P. teres* pathosystem. By employing the largest fungal mapping population for *P. teres* to date, this study successfully identified key QTL associated with virulence and confirm their correspondence to host susceptibility QTL. This study established the link between Ptt virulence on Skiff on Chr 3H and 6H, while Prior virulence on Chr 6H. These results suggest that possessing resistant 3H and 6H regions in barley breeding lines would provide resistance to Skiff and Prior virulent isolates. The identified Ptt candidate genes associated with virulence in Chr 5 for barley cultivar Prior would bring one step closer to confirm the virulence gene/s associated with *P. teres* barley pathosystem. Overall, this study provides valuable insights into the genetic basis of barley resistance to *P. teres* for breeders and lays the groundwork for future research aimed at developing disease-resistant barley cultivars.

## Author Contributions

B.D. and A.M. performed the QTL analysis. L.S provided the field data and recombinant inbred line populations. B.D. and S.B. performed the RNA experiment and extraction. B.D. performed the bioinformatics analyses including RNA sequencing analyses. P.B. and M.S. performed some parts of the phenotyping for Pop2 QTL analyses. B.D. prepared and revised the manuscript. S.B., L.S and A.M. reviewed the manuscript.

## Data Availability

Raw RNA sequencing data have been deposited to the NCBI under the submission code SUB15710218.

## Ethics, Consent to Participate, and Consent to Publish declarations

Not applicable

## Acknowledgement

This project was supported by the Australian Research Council Linkage Grant (ARC_LP220100084). Part of the funding for RNA sequencing was provided by the 2023 Early Career Seed Grant from the University of Southern Queensland. We acknowledge Judy McIlroy from the Hermitage Research Facility in Warwick, Australia, for supplying *Pyrenophora teres* t. *form* samples. We also thank our Master’s students Prashant Giri, Laxmi Kharel, and Laxmi Gautam Bhusal for their technical assistance.

